# Cell morphology best predicts tumorigenicity and metastasis *in vivo* across multiple TNBC cell lines of different metastatic potential

**DOI:** 10.1101/2023.06.14.544969

**Authors:** Sydney Conner, Justinne R. Guarin, Thanh T. Le, Jackson Fatherree, Charlotte Kelley, Samantha Payne, Ken Salhany, Rachel McGinn, Emily Henrich, Anna Yui, Savannah Parker, Deepti Srinivasan, Hanan Bloomer, Hannah Borges, Madeleine J. Oudin

## Abstract

**Background:** Metastasis is the leading cause of death in breast cancer patients. For metastasis to occur, tumor cells must invade locally, intravasate, and colonize distant tissues and organs, all steps that require tumor cell migration. The majority of studies on invasion and metastasis rely on human breast cancer cell lines. While it is known that these cells have different properties and abilities for growth and metastasis, the *in vitro* morphological, proliferative, migratory, and invasive behavior of these cell lines and their correlation to *in vivo* behavior is poorly understood. Thus, we sought to classify each cell line as poorly or highly metastatic by characterizing tumor growth and metastasis in a murine model of six commonly used human triple-negative breast cancer xenografts, as well as determine which in vitro assays commonly used to study cell motility best predict *in vivo* metastasis.

**Methods:** We evaluated the liver and lung metastasis of human TNBC cell lines MDA-MB-231, MDA-MB-468, BT549, Hs578T, BT20, and SUM159 in immunocompromised mice. We characterized each cell line’s cell morphology, proliferation, and motility in 2D and 3D to determine the variation in these parameters between cell lines.

**Results:** We identified MDA-MB-231, MDA-MB-468, and BT549 cells as highly tumorigenic and metastatic, Hs578T as poorly tumorigenic and metastatic, BT20 as intermediate tumorigenic with poor metastasis to the lungs but highly metastatic to the livers, and SUM159 as intermediate tumorigenic but poorly metastatic to the lungs and livers. We showed that metrics that characterize cell morphology are the most predictive of tumor growth and metastatic potential to the lungs and liver. Further, we found that no single *in vitro* motility assay in 2D or 3D significantly correlated with metastasis *in vivo*.

**Conclusions:** Our results provide an important resource for the TNBC research community, identifying the metastatic potential of 6 commonly used cell lines. Our findings also support the use of cell morphological analysis to investigate the metastatic potential and emphasize the need for multiple *in vitro* motility metrics using multiple cell lines to represent the heterogeneity of metastasis *in vivo*.

## Background

Metastasis, the dissemination of primary tumor cells to secondary organs in the body, is the leading cause of death in breast cancer patients. In the United States, it is estimated that 287,850 women will have been diagnosed with invasive breast cancer in 2022, and 43,250 women will have died of breast cancer [1]. The 5-year survival rate for localized breast cancer is 99%, but decreases to 80% once cells have reached the lymph nodes, and drops to 29% once metastases have formed in distant organs[1]. Triple-negative breast cancer (TNBC), which represents 10-15% of all breast cancers, is known to have higher rates of metastasis and disease recurrence with distant metastases [2], with even lower 5-year survival rates 8%-16% lower than non-TNBC [3]. TNBC lacks the expression of targetable receptors making it more difficult to treat [4] [5], with chemotherapy still being the standard of care for this subtype. Metastasis requires the local invasion of tumor cells into the surrounding environment, followed by entry into the lymphatics or vasculature and subsequent colonization of distant organs to form metastases [6], [7]. Metastasis occurs early on in cancer progression and significantly reduces survival in any cancer [8]. All steps of the metastatic cascade require tumor cell migration: tumor growth, local dissemination and intravasation, and colonization [9]–[11]. While mutations and epigenetic changes play a critical role in driving cancer progression, no single mutation renders cells metastatic or distinguishes a local from a metastatic tumor. Instead, metastasis is driven by multiple signaling pathways inside the cell in response to many different cues from the local tumor environment [12], [13]. Recent studies have suggested that chemotherapy drugs commonly used to inhibit tumor growth do not target invading cells and may promote local invasion of the surviving cells in breast tumors [14]–[16]. Eradicating metastatic breast cancer requires an in-depth understanding of the mechanisms that drive tumor cell migration.

*In vitro*, cell migration assays are commonly used to dissect the role of specific pathways in driving metastasis and identify drugs that could target metastatic disease by reducing cell migration. However, evaluating which cell migration assay is most informative of metastatic potential is an ongoing challenge. Many studies on cell motility have relied primarily on 2D assays where cells are seeded on a flat surface. Collective cell behaviors have been studied with the scratch wound assay, where cell migration is quantified by measuring wound closure over time, while single-cell motility behaviors can be assessed by 2D live imaging where both cell speed and persistence can be quantified [17]–[19]. Changes in cell morphology, driven by the cytoskeleton, are critical for cell migration. Cell morphology has been shown to be driven by distinct gene expression patterns and is a metric that can predict metastatic potential [20], [21]. We have also shown that cell morphology predicts the invasive behavior of TNBC cells in the context of extracellular matrix-driven migration [22]. 3D assays, where the cells are encapsulated in a 3D ECM scaffold, are known to better mimic the complex biochemical and physical properties of the native tissue microenvironment. Migration behavior in 2D versus 3D differs due to the more complex interactions between the microenvironment and the cytoskeleton in 3D [18], [23]. Spheroid assays, where cancer cells form aggregates encapsulated in a scaffold, have also been used to better mimic *in vivo* tumor dynamics [24], [25]. More complex assays involving microfluidic devices have also been developed and used to predict metastasis. Yankaskas et. al showed that the ability of cells to migrate through confined spaces, in combination with proliferation, could predict metastatic abilities in breast cancer [26]. Typically, a combination of these metrics is required by researchers to correlate or even predict *in vitro* cell behavior to *in vivo* response.

Most studies investigating factors regulating migration and metastasis rely on human breast cancer cell lines. Since the first breast cancer cell line BT20 was established in 1958 from the breast tumor of a 74-year-old female patient, numerous cell lines have been developed and used *in vitro* and *in vivo* [27], [28]. The most widely used TNBC cell line, MDA-MB-231 (231), has been cited 19,331 times on PubMed. Other commonly used cell lines are MDA-MB-468 (468), BT549, Hs578T, and SUM159 (Fig. S1). While it is known that these cell lines have different properties and abilities for growth and metastasis, these cell lines’ *in vitro* morphological, proliferative, migratory, and invasive behavior, and their correlation to *in vivo* behavior are poorly understood. Thus, we sought to classify each cell line as poorly or highly metastatic by characterizing tumor growth and metastasis to the lungs and livers in a murine model of six commonly used human triple-negative breast cancer xenografts 231, 468, BT549, Hs578T, BT20, and SUM159. We then characterized each cell line’s morphology, proliferation, and motility in 2D and 3D to determine the variations in these parameters between cell lines. We identify 231, 468, and BT549 cells as highly tumorigenic and metastatic, BT20 as intermediate tumorigenic, poorly metastasic to the lungs but highly metastatic to the livers, SUM159 as intermediate tumorigenic but poorly metastatic to the lungs and livers, and Hs578T as poorly tumorigenic and metastatic. Further, we show that metrics characterizing cell morphology are the most predictive of tumor growth and metastatic potential, but that no single *in vitro* motility assay significantly correlates with metastasis *in vivo*.

## Methods

### Animal Studies

The Tufts University Institutional Animal Care and Use Committee reviewed and approved all animal studies. Female NOD/SCID mice were obtained from The Jackson Laboratory (Bar Harbor, ME). At 6-8 weeks of age, female mice were injected with 2 million cells suspended in 20% collagen I (Corning, Rat Tail Collagen I) in PBS into the fourth left mammary fat pad using a 25G needle. Mice were monitored for tumor growth for 9 weeks. Tumor burden was monitored using digital calipers each week. After 9 weeks or until the maximum tumor burden of 1.5 cm^3^ was reached or significant ulceration, mice were euthanized by CO2 asphyxiation and cervical dislocation. Mammary tumors or fat pads, lungs, and livers were excised for further study.

### Histology

Mammary tumors, livers, and lung tissues were dissected, washed in PBS, and fixed for 24 hours in 4% paraformaldehyde in PBS before tissue processing in formalin. Tissues were then embedded in paraffin and sectioned into 10μm sections. For hematoxylin and eosin staining, sections were deparaffinized, hydrated, and stained using standard procedures with hematoxylin (GHS280; Sigma-Aldrich, St. Louis, MO), and counterstained with eosin (HT110180; Sigma-Aldrich, St. Louis, MO). Stained sections were mounted with toluene (SP15-500, Fisher Scientific, Hampton, NH). All slides were imaged using a Keyence BZ-X710 microscope (Keyence, Elmwood Park, NJ). Histologic analysis of hematoxylin and eosin (H&E) staining of these lungs and livers was quantified for metastases. Metastatic lesions were calculated by counting the number of metastases and the size of the metastasis using ImageJ.

### Cell Culture

MDA-MB-231, MDA-MB-468, BT20, BT549, Hs578T, and SUM159, cells were obtained from American Type Cell Collection (Manassas, VA). MDA-MB-231, MDA-MB-468, and BT20 were cultured in Dulbecco’s modified Eagle’s medium (DMEM) with 10% fetal bovine serum and 1% penicillin-streptomycin-glutamine. BT-549 and Hs578T were cultured in Dulbecco’s modified Eagle’s medium (DMEM) with 10% fetal bovine serum, 1% penicillin-streptomycin-glutamine, and 10μg/mL insulin. SUM159 were cultured in F12, 5% fetal bovine serum, 1% anti-anti, 5μg/mL insulin, 1 μg/mL hydrocortisone, and 20ng/mL epidermal growth factor (EGF). Cells were monitored for mycoplasma contamination every 2-3 months by polymerase chain reaction (PCR) using the Universal Mycoplasma Detection Kit (30-1012K; ATCC, Manassas, VA). All cells used in this study were mycoplasma negative.

### 2D cell Adhesion assay

Clear plastic plates coated with Collagen I protein were seeded with either MDA-MB-231, MDA-MB-468, BT20, BT549, Hs578T, and SUM159 cells (5,000 cells/well)and left to attach for 2 hrs. Then cells were fixed using 4% PFA for 10 min, permeabilized with 0.2% Triton-X-100, blocked with 3% bovine serum albumin (BSA), and stained to visualize their cytoskeleton (F-actin) with phalloidin and their nuclei with DAPI. Cells were imaged using a Keyence BZ-X710 microscope (Keyence, Elmwood Park, NJ) and the morphology of each cell was quantified using Cell Profiler. Cells were first identified from the nucleus, and the outline of each cell was determined from the cytoplasm staining. Cells at the edge of an image were discarded. 2D adhesion was quantified by parameters: area/cell (number of square μm in the cell cytoplasm), compactness (mean squared distance of the cell cytoplasm from the centroid divided by the area, where a filled circle has a value of 1, and an irregular shape has a value greater than 1), eccentricity (ratio of the distance between the foci of the ellipse and its major axis length, where a perfect circle has a value of 0, and more elongated cells have a value of 1), form factor (calculated as 4πArea/Perimeter^2^, where a perfect circle has a value of 1), perimeter, and solidity (proportion of pixels that are in the convex hull that are also in the cell cytoplasm, where a perfect circle has a value of 0). Data are the result of three independent experiments with three technical replicates per experiment.

### 2D Cell Proliferation

Cells were seeded on collagen I coated 96 well polystyrene plates (5,000 cells per well) and allowed to adhere for 24 or 72hrs. PrestoBlue™ Cell Viability Reagent (A13261; Invitrogen, Carlsbad, CA) was added to each well according to the manufacturer’s instructions and incubated for 30 min at 37°C. The absorbance of cells plated for 24hrs and 72hrs was measured on a plate reader and the fold change in absorbance was quantified between 24hrs and 72hrs.

### 2D Cell Migration

MDA-MB-231, MDA-MB-468, BT20, BT549, Hs578T, and SUM159 cells transfected with GFP were plated (5,000 cells/well) on plates coated with 1mg/ml collagen I protein for 1hr and allowed to adhere for 2 hours at 37°C before imaging. Cells were imaged every 20 minutes overnight for 16 hours using a Keyence BZ-X710 microscope (Keyence, Elmwood Park, NJ). Cells were then tracked using VW-9000 Video Editing/Analysis Software (Keyence, Elmwood Park, NJ), and migration speed and persistence were calculated using a custom MATLAB script vR2019a (MathWorks, Natick, MA).

### 3D Single Cell Invasion

MDA-MB-231, MDA-MB-468, BT20, BT549, Hs578T, or SUM159 cells transfected with GFP were resuspended in full media (400,000 cells/ml) with 1mg/mL collagen I (354236; Corning, Corning, NY), 10mM NaOH, 7.5% 10X DMEM, and 50% DMEM and plated in 96 well glass bottom plates. Plates were incubated at 37°C for 2 hours before imaging. Once ECM gelled, 50uL of culture media was added. Z-stack images of cells were captured every 20 minutes for 16 hours using the Keyence BZ-X710 microscope (Keyence, Elmwood Park, NJ). Cells were then tracked using VW-9000 Video Editing/Analysis Software (Keyence, Elmwood Park, NJ), and invasive speed and persistence were calculated using a custom MATLAB script vR2019a (MathWorks, Natick, MA).

### 3D Spheroid Invasion

MDA-MB-231, MDA-MB-468, BT20, BT549, Hs578t, and SUM159 cells transfected with GFP were seeded with full media in round-bottom, low-attachment 96-well plates (1,000 cells/well) and centrifuged at 3,000rpm for 3min to form spheroids. Spheroids were incubated at 37°C for three days. After 3 days spheroids were either grown in media or an ECM mixture of 1mg/mL collagen I (354236; Corning, Corning, NY), 10mM NaOH, 7.5% 10X DMEM, and 50% DMEM. Once ECM gelled, 50uL of culture media was added. Spheroids were imaged using a Z-stack on the day of ECM addition (day 0) and after 4 days of growth using a Keyence BZ-X710 microscope (Keyence, Elmwood Park, NJ). Spheroid invasion was quantified by measuring the distance cells on the periphery of the spheroid invade from the center and averaging for each spheroid. The fold change in spheroid invasion distance was then quantified relative to the average invasion distance of no ECM control spheroids for each cell line. The data presented are the result of three independent experiments with three technical replicates per experiment.

### Statistical analysis

GraphPad Prism v9.1.0 was used for the generation of graphs and statistical analysis. Significance was determined using a one-way ANOVA with Dunnett’s multiple comparisons test unless otherwise stated. MATLAB was used to determine linear relationships and correlation. For all graphs, we performed statistical analysis in comparison to 231, given that this is the most commonly used cell line in the literature.

## Results

### Characterization of Tumor Growth and Metastasis of cell line xenografts *in vivo*

There are currently no studies that directly compare the metastatic ability of multiple TNBC cell lines, with the same murine xenograft model and experimental parameters. We first directly compared the growth and metastasis of six human TNBC cell lines commonly used to study mechanisms of TNBC metastasis. We used 231, 468, BT549, Hs578T, and SUM159 cell lines, which were isolated from patients with different racial backgrounds and tumor types and which have different mutational profiles and properties (Fig 1A). We injected the same concentration of cells in a collagen I solution into the 4^th^ left mammary fat pad of female NOD-SCID mice. Tumors were left to grow for 9 weeks or until ulcers began to form. We found that 468 tumors grew the fastest, followed by 231 and BT549 tumors which had similar sizes. BT20 and SUM159 xenografts grew the smallest tumors, and at 8 weeks the SUM159 tumors had ulcerated and had to be sacrificed. The Hs578T xenografts did not grow palpable tumors in this time frame (Fig 1B, C). These data demonstrate that 231, 468, and BT549 cell lines are highly tumorigenic, BT20 and SUM159 are intermediate tumorigenic, and Hs578T are poorly tumorigenic since they did not form palpable tumors.

**Figure 1.**
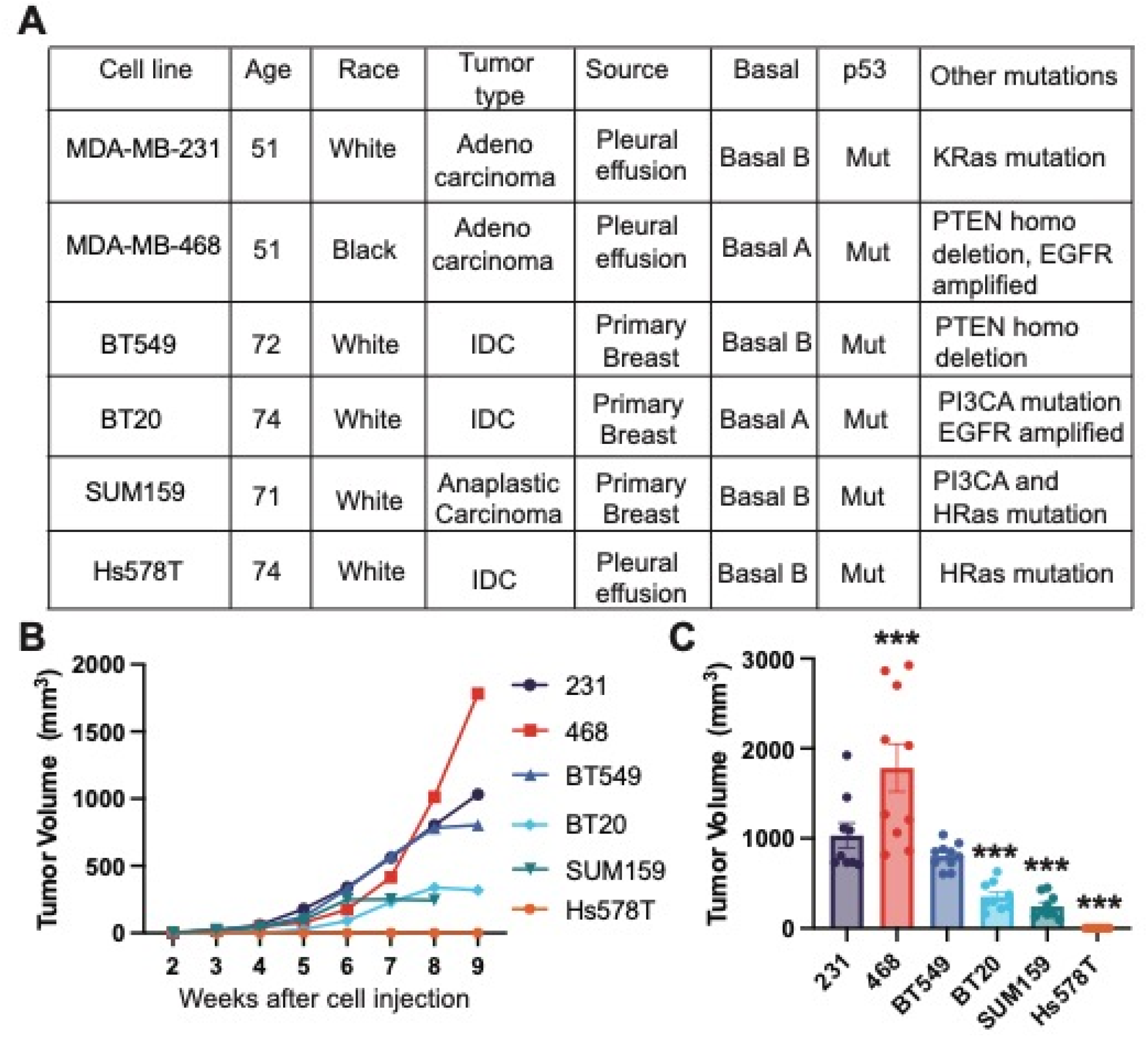
Tumor growth of human TNBC cell xenografts implanted in immunocompromised female NOD-SCID mice. (A) Characteristics of six human TNBC cell lines used in the study. M, mutant; WT, wild type; Mes, Mesenchymal; Bas, Basal; IDC, invasive ductal carcinoma; (B) Tumor volume over time for each cell line 231, 468, BT549, Hs578T, BT20, and SUM159. (C) Tumor volume at the terminal endpoint. Data are shown as mean with SEM, n = 8-10 animals/group. Significance relative to 231 was determined using a one-way ANOVA with Dunnett’s multiple comparisons tests, *p<0.05, ** p<0.01, *** p<0.005.

Next, we examined lung and liver metastases, by quantifying the number of metastases found in each organ, the average size of each metastatic lesion, and the metastatic index, which is the number of metastases relative to primary tumor size. The 231, 468, and BT549 cell lines had the most lung metastases and similar metastatic index. The BT20, Hs578T, and SUM159 cells had very low numbers of lung metastases, with very low metastatic index (Fig 2A-C). The size of lung metastases varied between cell lines with 231s having significantly larger metastases than BT549 and 468 (Fig 2D). The lung metastatic lesions in BT20, Hs578T, and SUM159 were very small (Fig 2D). We then examined the relationship between primary tumor size, number of lung metastases, and size of lung metastases. For all cell lines, there was a significant correlation between primary tumor volume and the number of metastatic lesions in the lung (Fig S2A, B). There was no significant correlation between the number and sizes of lung metastases (Fig S2C, D) or between the size of the primary tumor and the size of the metastases in the lungs (Fig S2E, F).

**Figure 2.**
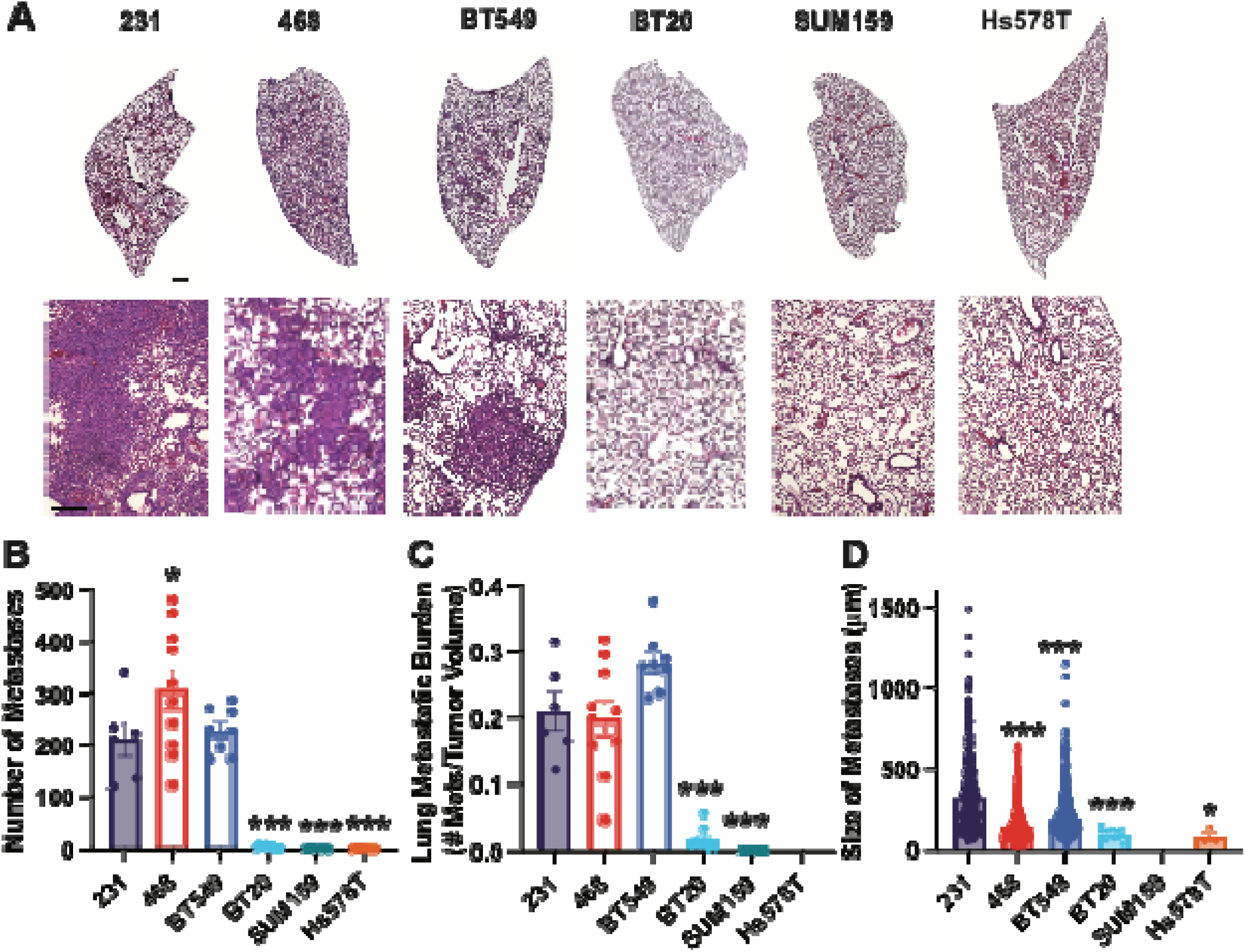
TNBC cells have different metastatic potentials to the lung. (A) Representative lung tissue sections stained with H&E. Scale bar = 500μm Inset scale bar = 500μm. (B) The number of metastases per lung for each cell line and (C) Metastatic burden in lungs (ratio of the number of metastases in the lungs to the primary tumor volume) (D) Average size of each lung metastasis. Data are shown as mean with SEM n = 8-10 animals/group. Significance relative to 231 was determined using a one-way ANOVA with Dunnett’s multiple comparisons tests, *p<0.05, ** p<0.01, *** p<0.005.

The BT549, 231, and 468 cell lines developed the most liver metastases, with similar levels of metastatic index and size of metastases (Fig 3A-D). The BT20 xenografts resulted in a high number of liver metastases, which given their small tumors, led to a high liver metastatic index for this cell line, significant relative to 231 cells (Fig 3A-D). Interestingly, for the BT20 cell line while the number of liver metastases was not significantly different from the 231 cell line the size of the metastases was significantly smaller (Fig 3A-D). SUM159 and Hs578T xenografts both developed significantly fewer liver metastases than 231s (Fig 3A-D), which were also smaller. We then examined the relationship between primary tumor size, number of liver metastases, and size of liver metastases. In the livers, there were differences between the highly metastatic and poorly metastatic cell lines. There was a significant correlation between primary tumor volume and the number of metastatic lesions in the liver, and the number and size of liver metastases but only for the least metastatic cell lines (BT20, Hs578T, and SUM159) (Fig S2G-J). For the highly metastatic cell lines (231, 468, and BT549), there was a negative significant correlation between primary tumor size and liver metastases size (Fig S2E, F).

**Figure 3.**
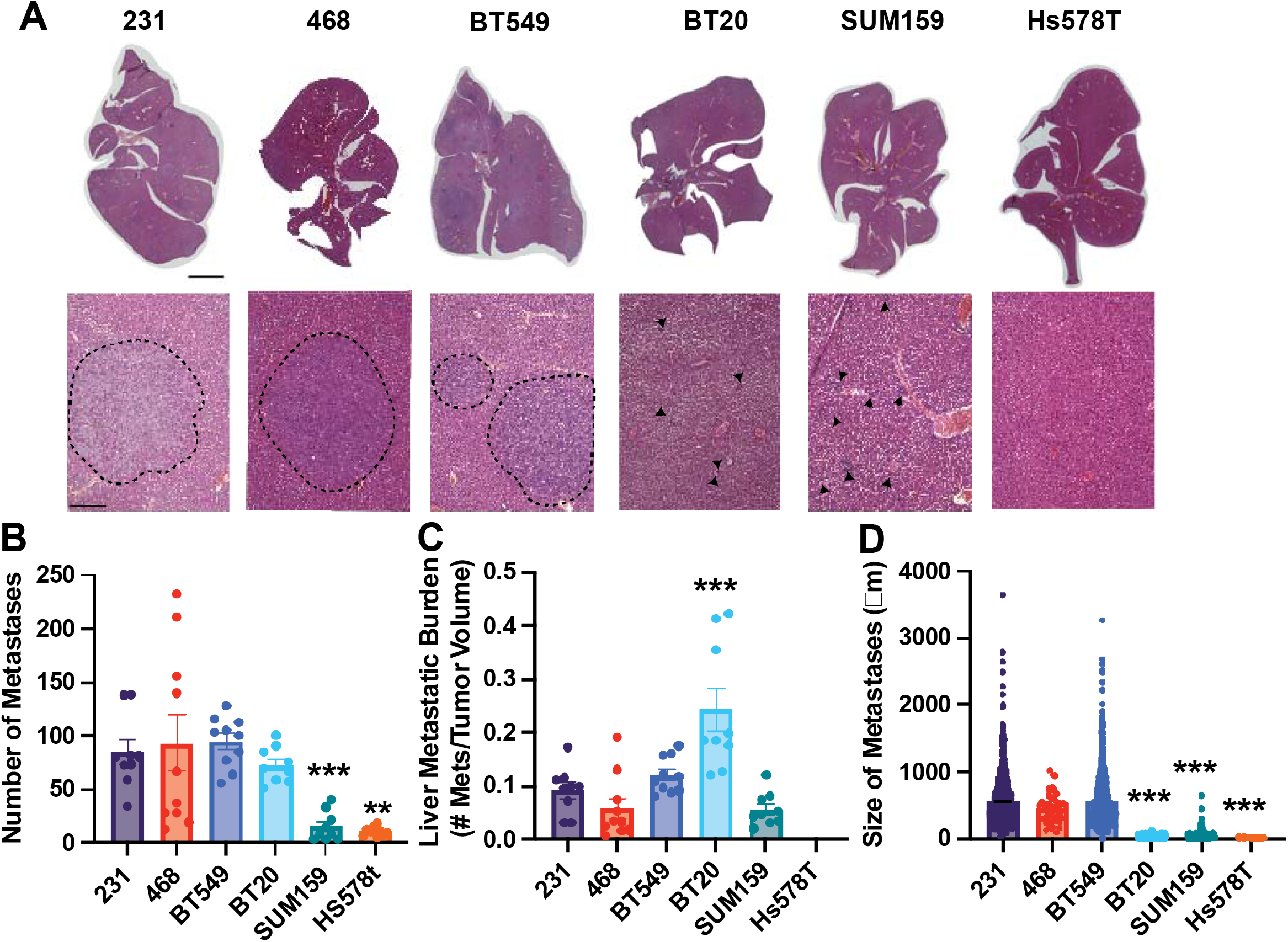
TNBC cells have different metastatic potentials to the liver. (A) Representative liver tissue sections stained with H&E. Scale bar is 1mm; inset scale bar is 500μm dashed lines represent metastasis boundary, black arrows identify smaller metastases. (B) The number of metastases per liver for each cell line and (C) Metastatic burden in the liver (ratio of the number of metastases in the lungs to the primary tumor volume) (D) Average size of each liver metastasis. Data are shown as mean with SEM n = 8-10 animals/group. Significance relative to 231 was determined using a one-way ANOVA with Dunnett’s multiple comparisons tests, *p<0.05, ** p<0.01, *** p<0.005.

These data characterize the tumorigenicity and metastatic potential of six human TNBC cell lines, demonstrating that 231, 468, and BT549 cell lines are highly tumorigenic, BT20 and SUM159 are intermediate tumorigenic, and Hs578T are poorly tumorigenic since they did not form palpable tumors. These data characterize 231, 468, and BT549 cell lines as highly metastatic to both the lungs and the livers, BT20 as poorly metastatic to the lungs but metastatic to the liver, and SUM159 and Hs578Ts as poorly metastatic to both the lungs and the liver with slight dissemination to the liver.

### TNBC cell lines have distinct differences in morphological characteristics

We next investigated the *in vitro* characteristics of six TNBC cells. It has been previously shown that features of cell morphology correlate to cell motility and metastatic potential [20], [21]. 2D cell adhesion assays are commonly used to investigate cell phenotypic behavior because they are relatively inexpensive and have low equipment requirements. Cells were left to adhere to Collagen I, the most abundant ECM protein in breast tissue, and their morphology was analyzed: size (cell area and perimeter), irregularity (solidity and form factor), and elongation (eccentricity and compactness) were quantified (Fig 4A). The 468 cells were the smallest, as measured by cell area and perimeter (Fig 4B, C, S3A), they were also less irregular than 231 cells indicating fewer protrusions measured by solidity and form factor, but also rounder and less elongated, measured by eccentricity and compactness. (Fig 4D, E, S3B, C). BT549 and BT20 cells were slightly larger than 231s, exhibited a more rounded morphology, and were less irregular, with fewer protrusions (Fig 4B-E, S3 A-C). We found that Hs578Ts were the largest cells with more irregular shapes indicating more protrusions and were also rounder and less elongated (Fig 4B-E, S3A-C). SUM159 cells were not as large but were similarly more protrusive and more elongated (Fig 4C-E S3A-C). SUM159s were very irregular, with more protrusions, and were the most elongated (Fig 4C-E S3A-C).

**Figure 4.**
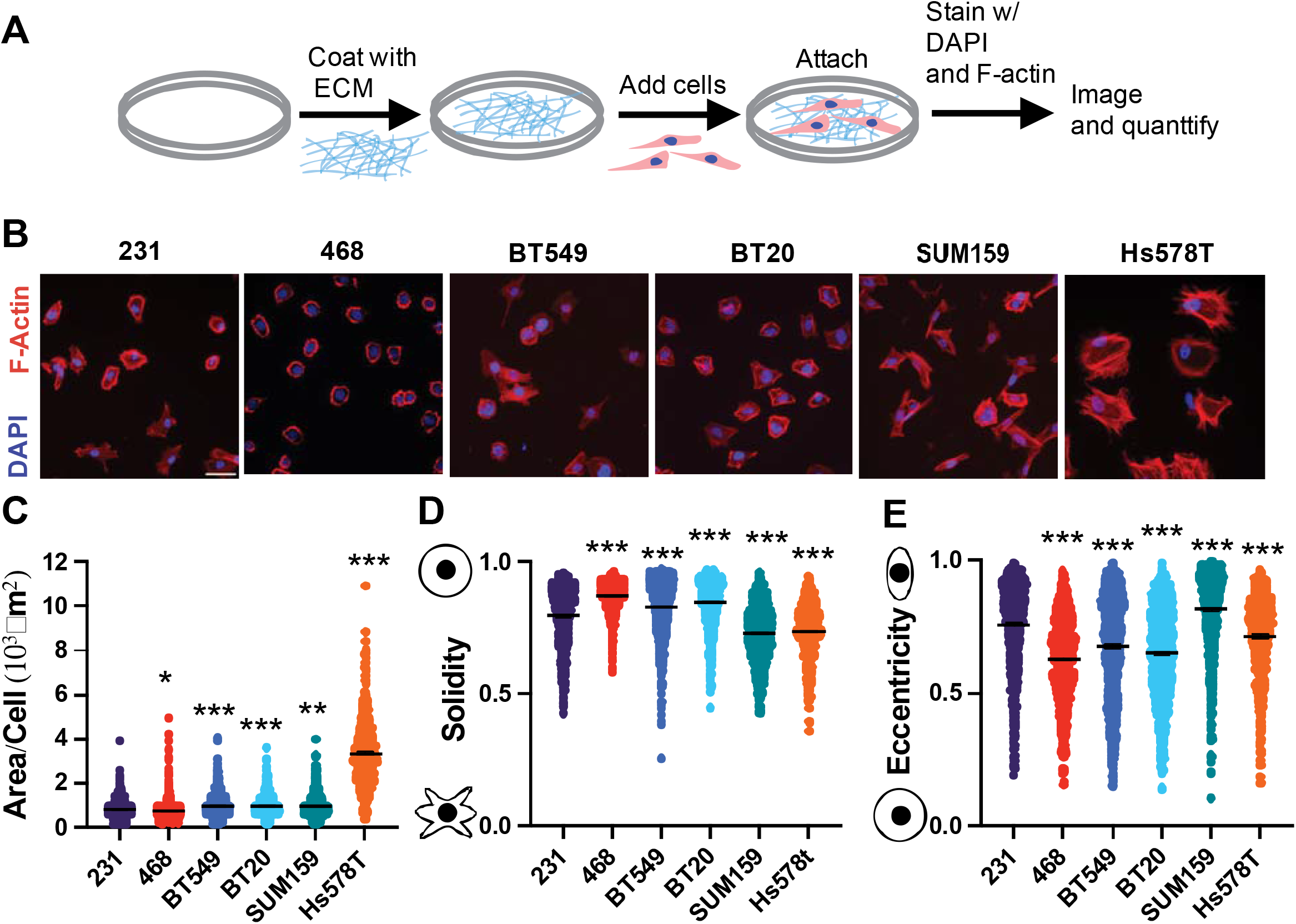
Characterization of TNBC cell line morphology. (A) Schematic depicting experimental procedure. (B) Representative images of 231, BT549, Hs578T, 468, BT20, and SUM159 cell lines plated on Collagen I ECM fixed and stained for nuclei (Blue) and F-actin (red). Scale bar = 50μm. Quantification of cell shape parameters for each cell line 231 (n=679 cells), BT549 (n=799 cells), Hs578T (n=538 cells), 468 (n=818 cells), BT20 (n=882 cells) and SUM159 (n=892 cells) (C) Area/Cell (D) Solidity (E) Eccentricity Data show mean ± SEM. Significance was determined using a one-way ANOVA with Dunnett’s multiple comparisons test compared to 231 cell line *p<0.05, ** p<0.01, *** p<0.005. Each data point represents a single cell.

Principal component analysis (PCA) was used to reduce the dimensionality of the adhesion data set, incorporating all six cell adhesion metrics into two principal components (PCs). The loading scores represent how the independent variables project on each principal component. The loadings scores represent variations in cell morphology metrics. As expected, perimeter and area, solidity and form factor, and eccentricity and compactness clustered together (Fig S3D). The scores in the principal component space represent the variation in cell morphology among cell lines. The highly tumorigenic and metastatic cell lines, 231, 468, and BT549 clustered together, while the poorly tumorigenic and metastatic SUM159s, Hs578Ts, and BT20s lines appeared more distant from this cluster indicating that highly tumorigenic and metastatic cell lines share more similar morphological characteristics than poorly tumorigenic and metastatic cell lines (Fig S3E).

### Proliferation and 2D cell motility

Cell proliferation is critical to support both primary and secondary tumor growth. We quantified the proliferation rate of each cell line by measuring the change in the metabolic activity over 48hrs (Fig 5A). BT549 and SUM159 cell lines were significantly more proliferative than 231s (Fig 5B). 2D cell migration is a commonly used metric to determine metastatic potential. Cells were seeded on Collagen I and imaged to quantify 2D cell migration speed, which measures how fast the cell is moving over the distance traveled, and persistence which measures the Euclidean distance between the start and finish of the cell’s path over the total distance traveled. SUM159, 231, and 468 cells migrated the fastest in 2D and were not significantly different from each other. Hs578T, BT20, and BT549 cells all migrated significantly slower than 231 cells (Fig 5D). 231, 468 and BT20 cells had the lowest 2D persistence with BT594s, Hs578Ts and SUM159s all having significantly higher persistence than 231s respectively (Fig 5E,F).

**Figure 5.**
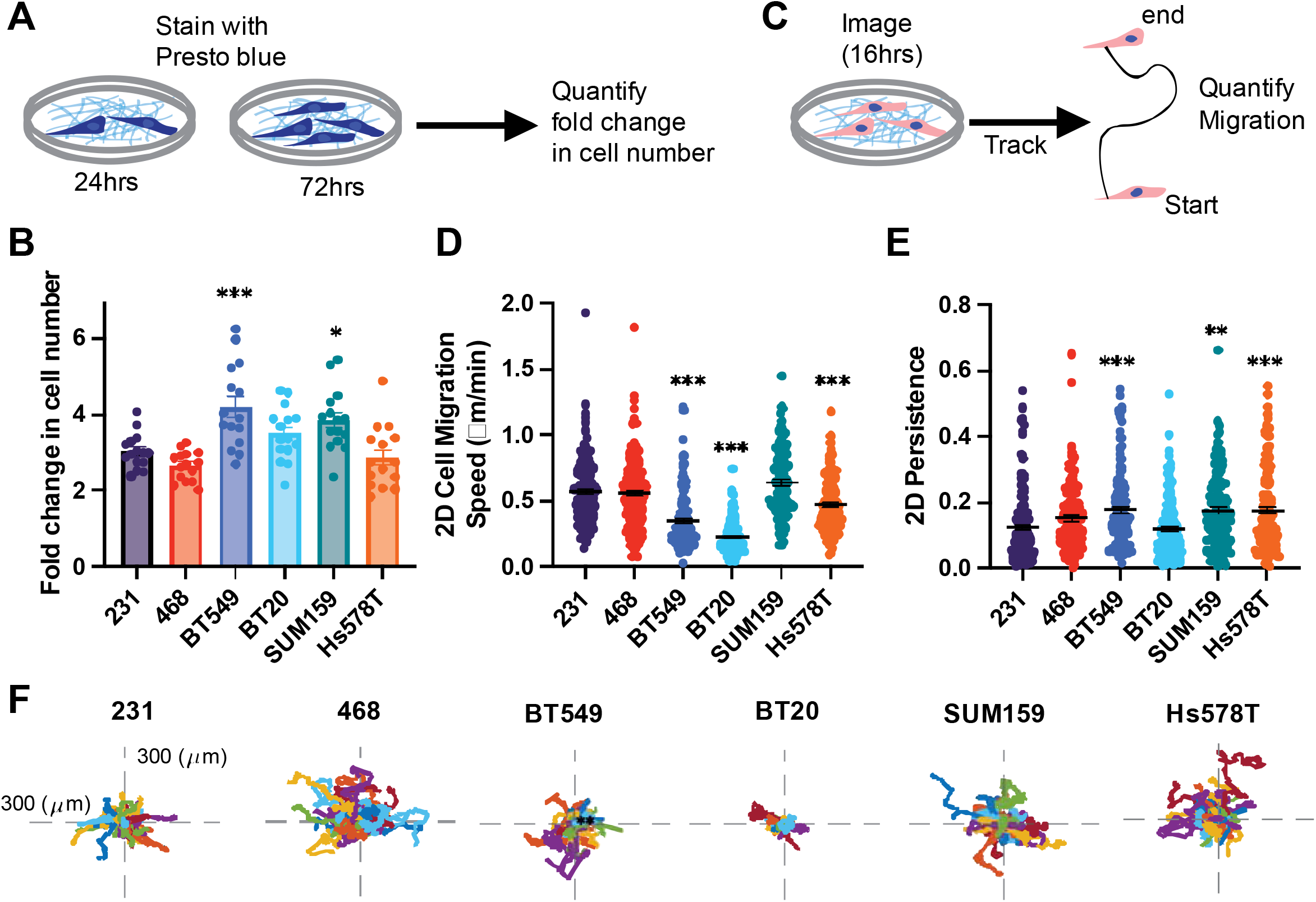
Cell line-specific proliferation, 2D migration, and persistence. (A) Schematic depiction of presto blue proliferation assay. (B) Schematic depiction of 2D migration assay. (C) The relative proliferation of cell lines (D) 2D cell migration and (E) 2D persistence of cell lines seeded on collagen 1 coated glass bottom plates. (F) representative rose plots of the migration of cells on collagen I ECM-coated glass coverslips. Data show mean ± SEM. Significance was determined using a one-way ANOVA with Dunnett’s multiple comparisons test compared to the 231 cell line *p<0.05, ** p<0.01, *** p<0.005. Each data point represents a single cell.

### 3D methods to quantify cell invasion

To better recapitulate the native breast microenvironment, 3D assays where breast cancer cells are encapsulated in ECM, are used to mimic the 3D *in vivo* environment. We first investigated 3D invasion in a single-cell invasion assay, where each cell line was encapsulated in Collagen I ECM and plated in a glass bottom dish then imaged overnight to track cell invasion (Fig 6A). The 468, SUM 159, and Hs578T cells invaded significantly faster than the 231 cells, and BT549 and BT20 cells invaded significantly slower than the 231 cells (Fig 6B). Interestingly, only SUM159 and BT20 cells were significantly more persistent than 231 cells (Fig 6 C, D).

**Figure 6.**
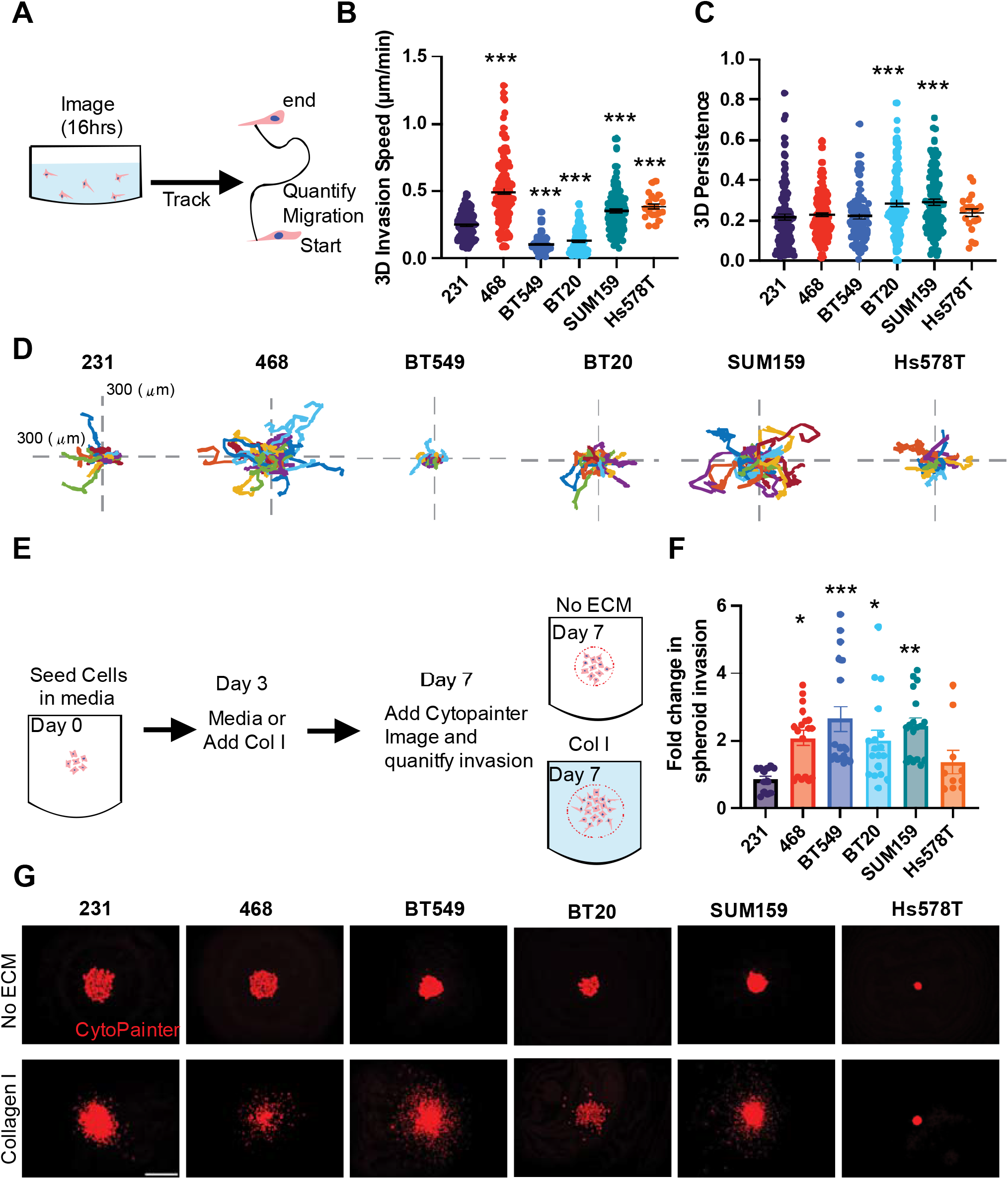
3D single cell invasion and spheroid invasion in Collagen I ECM. (A) schematic depicting experimental set up of 3D invasion assay. (B) 3D invasion of breast cancer cell lines invading through collagen I ECM (C) 3D persistence of cells invading through collagen I ECM (D) representative rose plots of the invasion of cells through the collagen I ECM. (E) schematic depiction of spheroid invasion assay. (F) Fold change in spheroid invasion into collagen I hydrogels over invasion into media (G) Representative spheroid images of 231, BT549, Hs578T, and SUM159 spheroids cultured in media (upper) or spheroids cultured in Collagen I solution visualized with cytopainter (red). Scale bar = 500μm. Data show mean ± SEM. Significance was determined using a one-way ANOVA with Dunnett’s multiple comparisons test compared to 231 cell line *p<0.05, ** p<0.01, *** p<0.005, ****P=<0.0001. Each data point represents a single cell.

We then used a 3D spheroid assay, where the cell lines were seeded in low attachment U-bottom dishes, to form a densely packed spheroid. After growing for 3 days either Collagen I was added, or the spheroids were left in media and grown for an additional 4 days. Growth out of spheroids relies on tumor cell proliferation and invasion. We then measured the fold change in the spheroid outer area in the Collagen I group relative to media only after 7 days (Fig 6E). 468, BT20, BT549, and SUM159 cells had a significantly higher fold change in spheroid invasion compared to 231s (Fig 6F-G).

PCA was then used to reduce the dimensionality of the data set to compare 2D migration and persistence and 3D invasion and persistence. The scores of each cell line demonstrate that highly tumorigenic and metastatic cell lines, 231, 468, and BT549 clustered together and the poorly tumorigenic and metastatic SUM159s, Hs578Ts, and BT20s clustered together indicating that highly and poorly tumorigenic and metastatic cell lines have similar motility characteristics (Fig S4A). Interestingly, the loadings scores represent variations in the motility metrics with 2D migration and persistence and 3D invasion cluster together while 3D persistence is by itself.

### Cell shape is most correlated with metastatic potential

Our last goal was to determine which assays and their metrics, cell morphology, proliferation, and 2D or 3D motility, best correlate with *in vivo* tumor growth and metastasis. To do this we used pairwise comparisons to systematically investigate each *in vitro* metric in relation to *in vivo* response for all cell lines. First, we determined if there was a linear relationship between each *in vitro* metric and *in vivo* response by calculating the R^2^ which measures the proportion of variation in the *in vivo* response that is attributed to each *in vitro* metric, and the Pearson correlation which measures the strength of the linear relationship. There were distinct differences in the linear relationship between *in vitro* metrics and *in vivo* behaviors. Interestingly, we found that solidity and form factor, which quantify how irregular or protrusive a cell is, had a linear relationship with tumor volume (R^2^>0.5) (Fig 7A, S5). Only the persistence of 3D single cell invasion had a linear relationship to the number of lung metastases (R^2^>0.5) (Fig 7A, S5). Cell size and cell irregularity, all had linear relationships to the number of liver metastases with solidity and form factor having a strong linear relationship (R^2^>0.7 and p<0.05) (Fig 7A S5). Next, we used Spearman’s correlation to investigate the correlation of each assay to *in vivo* response. We found that area, perimeter, and 3D persistence were negatively correlated with tumor volume, while solidity and form factor were positively correlated (Spearman Coefficient > ± 0.6). Area, perimeter, eccentricity, compactness, and 3D persistence demonstrated a negative correlation with the number of lung metastasis while solidity and form factor demonstrated a positive correlation. Perimeter and 3D persistence negatively correlated to the number of liver metastases, while solidity and form factor positively correlated (Fig 7B).

**Figure 7.**
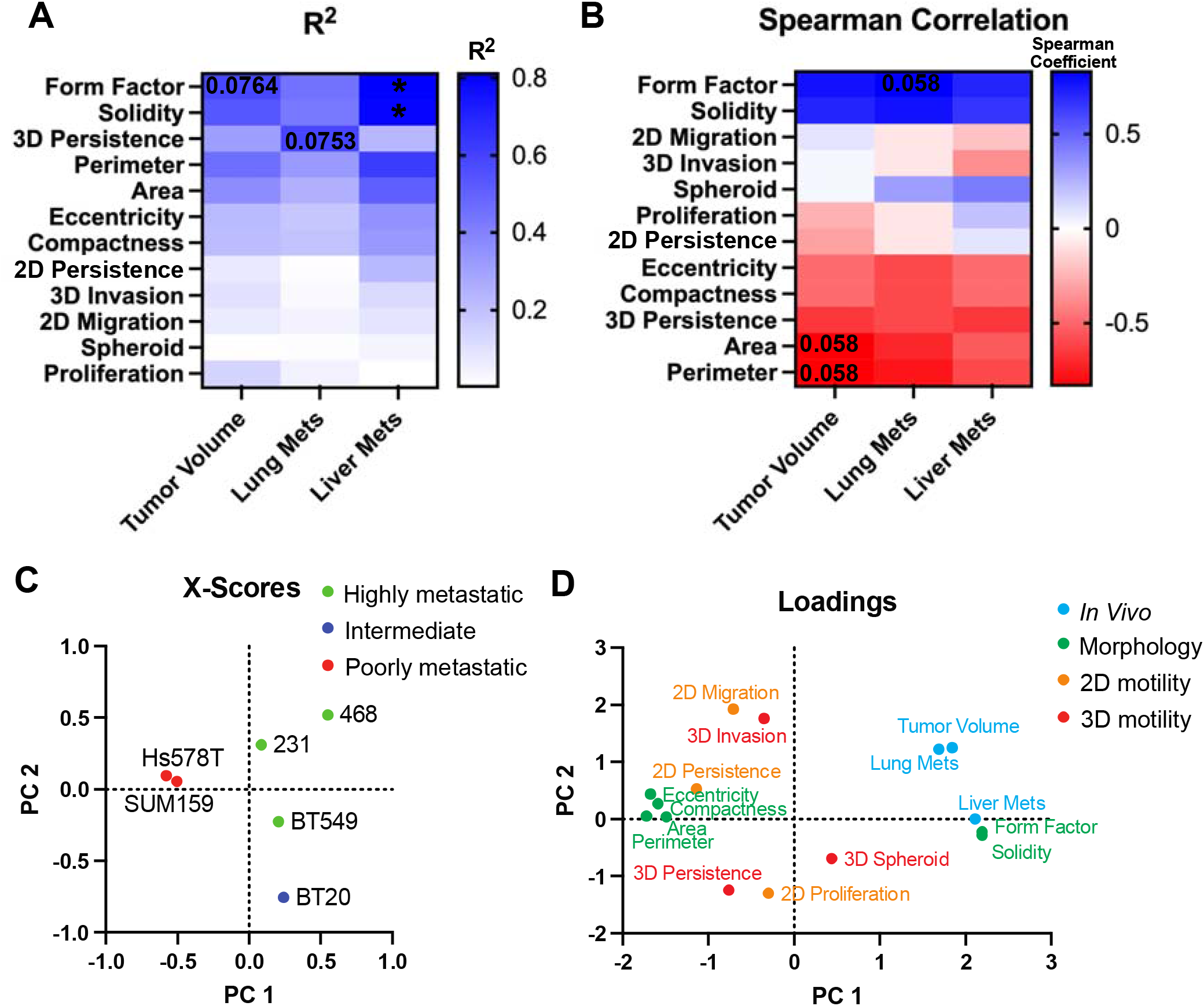
Distinct differences in cell line-specific behavior *in vitro* and *in vivo*. (A) pairwise analysis of the strength of the linear relationship (R^2^) between *in vitro* metrics and *in vivo* behavior. Numbers represent p-value and colors represent R^2^ value where blue is high and white is low indicated by gradient (B) Spearman correlation between *in vitro* metrics and *in vivo* behavior. Numbers represent p-value and colors represent the Spearman correlation coefficient where blue is a high positive correlation, white is no correlation and red is a negative correlation indicated by gradient (C) X-scores plot of the PLS model (D) PLS loadings for adhesion metrics, proliferation, 2D and 3D motility, and *in vivo* tumor volume, lung and liver metastases.

To visualize the relationship between the cell lines and assays we used partial least squares to calculate the scores and loadings which capture the covariance between the independent and dependent variables [29]. The scores plot describes how each cell line projects on principal components 1 (PC1) and PC2. Highly metastatic cell lines 231, 468, and BT549 clustered together, and poorly metastatic cell lines SUM159s and Hs578T clustered together. BT20s, which were intermediate tumorigenic but poorly metastatic to the lungs but metastatic to the livers, stood alone (Fig 7C). The loading scores represent how the independent variables and dependent *in vivo* responses project on each principal component. As expected, cell shape parameters related to size, elongation, and irregularity clustered together. Other cluster groups include lung metastasis and tumor volume; liver metastasis, form factor, and solidity; eccentricity, compactness, area, perimeter, and 2D persistence; 2D migration and 3D invasion; 2D, 3D persistence and proliferation (Fig 7D). This analysis demonstrates that there were distinct differences in cell line-specific behaviors *in vitro* and *in vivo* and that parameters that measure cell shape and 3D persistence are most correlated with *in vivo* behaviors.

## Discussion

Established human breast cancer cell lines and murine xenograft models are powerful tools in breast cancer research that provide a comprehensive picture of disease progression and the metastatic cascade. However, there is significant heterogeneity within cell lines in terms of their behaviors both *in vitro* and *in vivo. In vitro*, methods are preferred for investigating mechanisms, identifying therapeutic targets, and testing hypotheses before moving to costly and time-consuming *in vivo* models. As such, morphology, proliferation, migration, and invasion assays are often employed to characterize a cancer cell’s potential to grow and metastasize. Yet how metrics quantified in these assays correlate with *in vivo* behaviors such as tumor growth and metastasis is poorly understood. In this study, we characterized the metastatic potential of 6 commonly used human TNBC cell lines both *in vitro* and *in vivo*. We generated murine xenografts to quantity tumor growth, and lung and liver metastasis and performed several *in vitro* assays used to study cell behaviors such as morphology, proliferation, migration, and invasion, which are all mechanisms critical for tumor growth and metastasis. We then conducted analyses to better understand how the *in vitro* morphological, proliferative, migratory, and invasive behavior of TNBC cell lines correlate to their *in vivo* behavior to determine which assays best predict tumor growth and metastasis. We identified 231, 468, and BT549 cell lines as highly tumorigenic, BT20 and SUM159 as intermediate tumorigenic, and Hs578T as poorly tumorigenic. We identified 231, 468, and BT549 cell lines as highly metastatic to both the lungs and the livers, BT20 as poorly metastatic to the lungs but highly metastatic to the liver, and SUM159 and Hs578Ts as poorly metastatic to both the lungs and the liver. We found a positive correlation between primary tumor volume and both liver and lung metastasis for most of the cell lines. This is in line with patient data, where multiple studies have shown that primary tumor volume is an independent prognostic factor for distant metastasis and poor disease survival [30], [31]. We determined that parameters measuring cell shape and 3D persistence are most correlated with *in vivo* behaviors.

When studying the role of a particular step in metastasis or the effect of a drug or intervention on metastasis *in vivo*, it is important to consider the metastatic potential of the specific cell lines being used, as well as using multiple cell lines of different metastatic potential to cover a range of human phenotypes to ensure reproducible findings. One common approach involves using TNBC cells as a highly metastatic model, ER-positive MCF7 cells as a poorly metastatic model and healthy MCF10A as a control [32]. However, it is well known that the biology of TNBC and hormone-positive cancers are unique and that using different subtypes to study metastasis may yield less conclusive results. Another strategy includes using metastatic derivatives of cell lines that were generated after serial selection and *in vivo* passaging, such as the LM2 cell line derived from MDA-MB-231, which has a high propensity to the lung [33]. While these cell lines have significantly contributed to a better understanding of the mechanisms by which cells metastasize to the lung, they originate from the same MDA-MB-231 cell line and have been artificially selected for use in mouse models. Therefore, they do not necessarily represent a broad set of patient cell lines exhibiting different metastatic behaviors. The use of multiple TNBC cell lines of different metastatic potentials within a study helps to capture this population. However, prior to our study, there was no direct comparison and objective quantification of the metastatic potential of each of these cell lines. We focused on six of the most published TNBC cell lines, injected with the same number of cells in the same conditions and mouse model to determine their metastatic potential relative to each other. Our results demonstrate that 231, 468, and BT549 cell lines are highly tumorigenic and metastatic, BT20 is intermediate tumorigenic with poor metastasis to the lungs but metastatic to the livers, SUM159 is intermediate tumorigenic but poorly metastatic to the lungs and livers, and Hs578T is both poorly tumorigenic and metastatic. We believe these data can help rectify the existing discrepancies in the literature, where different cell lines are assigned metastatic potential without sufficient supporting evidence.

Several previous studies have made inaccurate classifications regarding the metastatic potential of these cell lines. In one study, 468 cells were assigned low metastatic potential, while Hs578T, BT549, SUM159, and 231 cells were assigned high metastatic potential [26]. In another study, 231 and SUM159 were labeled as highly metastatic [32]. We believe these misclassifications are likely due to the lack of a comprehensive dataset that directly compares these cell lines in vivo metastatic potential under consistent experimental conditions. Lastly, the recent Metmap study characterized the metastatic capability of 21 basal-like breast cancer cell lines; however, cells were injected intracardially, quantifying exclusively experimental metastasis and not spontaneous metastasis from the primary tumor site [34]. Experimental metastasis relies more on survival in the bloodstream, extravasation, and metastatic outgrowth, while spontaneous metastasis from the mammary gland additionally requires local invasion and intravasation [35]. Interestingly, we saw some similarities and striking differences between the metastatic potential of human TNBC cell lines based on the model used. 231 cells efficiently metastasize to the lung and liver at high rates in both models and BT20s metastasized to the livers but not the lungs from the intracardiac injection, similar to the spontaneous model. However, 468 cells showed poor metastatic potential from the intracardiac injection with most going to the liver, while in our model, these cells metastasized very efficiently to the lung and liver from the mammary gland. With the intracardiac injection, BT549 cells showed no metastasis to any organ, whereas they exhibited robust metastasis to both the lung and liver from the mammary gland in our study, comparable to 231 cells [34]. Overall, our data provide a clear characterization of the metastatic potential of six human TNBC cell lines in a spontaneous metastasis model from the mammary gland. Further, our study demonstrates that the metastatic potential from the primary tumor may be very different from that in an intracardiac injection metastasis model.

Our data suggest that factors characterizing cell morphology are highly predictive of tumor growth and metastatic potential to the lungs and liver. Interestingly, in the 2D adhesion assay the smaller, rounder, and less irregular cell lines 231, BT549, and 468 had larger tumors and more lung and liver metastases while the larger and more irregular cells Hs578Ts and SUM159s had smaller tumors and fewer metastases. These data align with many existing studies that highlight the importance of cell morphology in predicting metastatic behaviors. Indeed, single-cell clones derived from the 231 cell line demonstrated persistent morphological heterogeneity, which was correlated with metastatic potential *in vivo* [20]. This study focused less on individual shape factors but instead incorporated data from 216 features characterizing cell and nuclear morphology. Zaritsky *et al*. used a generative neural network in combination with supervised machine learning to predict the metastatic potential of patient-derived melanoma xenograft and found that pseudopodal extensions were the hallmark properties of metastatic cells [36]. Using murine and human osteosarcoma cells, another study found that cell shape could be used to distinguish the less metastatic cells from the more metastatic cells, with the more metastatic cell lines also smaller, rounder, and less irregular [37]. Many factors can impact cell morphology, and it is important to be mindful of the substrates used when studying the morphology of cultured cells. We have shown that different ECM proteins’ impact on morphology can be used to predict 3D invasion [22]. In our current study, we used Collagen I as a substrate, as it is the most abundant ECM protein in breast tissues [38], [39] In conjunction with existing studies, our data further support the use of cell morphological measurements to dissect the role of individual genes and pathways in metastasis.

Lastly, our study raises some questions about which *in vitro* cell motility assay and metrics are best to predict metastasis *in vivo*. Our data show that no single *in vitro* motility assay significantly correlates with metastasis *in vivo*. The metric that is most closely associated with lung metastasis was the 3D persistence of single cells invading in collagen I gel. It is possible that the substrates and local environment used in our studies are not representative enough of the complex ECM found in tissues. Recent advances using decellularized whole ECM scaffolds [38], cell-derived ECM [40], and co-culture systems [41] have sought to better mimic the tumor microenvironment. However, these complex systems make it difficult to parse out the contributing factors in cell behavior compared to simpler systems. In addition, the same cell line once cultured in different labs or under different conditions can evolve into distinct groups [28]. It is also possible that other components of the tumor microenvironment such as stiffness, and factors secreted by resident local and stromal cells, which are absent in these *in vitro* assays, may also impact the results. Interestingly, when metrics were combined, cell shape metrics or 2D motility and 3D invasion, we could cluster the cell lines of different metastatic potential by PCA (Fig S3D-E, S4). These data suggest that using multiple *in vitro* motility assays can offer a more accurate prediction of TNBC *in vivo* metastasis.

## Conclusions

Overall, we hope this work will provide a helpful resource for the TNBC research community, showing the first direct comparison of metastatic potential among commonly used TNBC cell lines. We encourage researchers to use multiple TNBC cell lines of different metastatic potential to better represent the heterogeneity of the human TNBC population, as well the use of multiple *in vitro* cell shape and motility assays to study the role of cell-intrinsic and cell-extrinsic features in driving metastasis in TNBC.

## List of abbreviations

ECM: Extracellular Matrix
PCA: Principal Component Analysis
PLSR: Partial Least Squares Regression
TNBC: triple-negative breast cancer

## Funding

This work was supported by:

National Institutes of Health [R00-CA207866, DP2CA271387 and R01CA255742 to M.J.O.],

Tufts University [Start-up funds from the School of Engineering to M.J.O.]

## Competing Interest Statement

Authors declare they have no competing interests

## Author contributions

All authors contributed to data generation and analysis, SJC and MJO wrote the manuscript.

The datasets used and/or analysed during the current study are available from the corresponding author on reasonable request.

## Supplemental

**Supplemental Figure 1.**
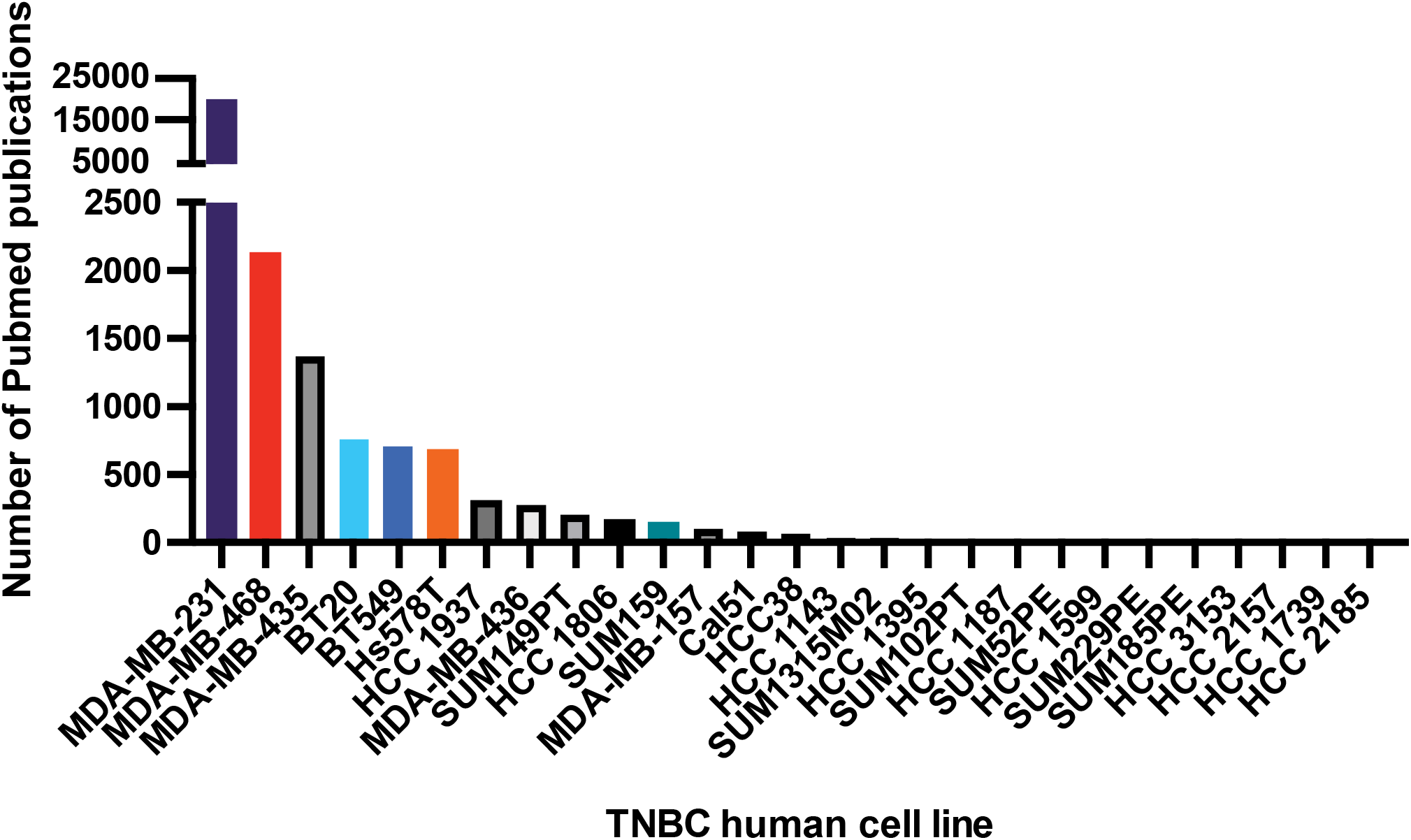
Prevalence of TNBC cell lines in literature.

**Supplemental Figure 2.**
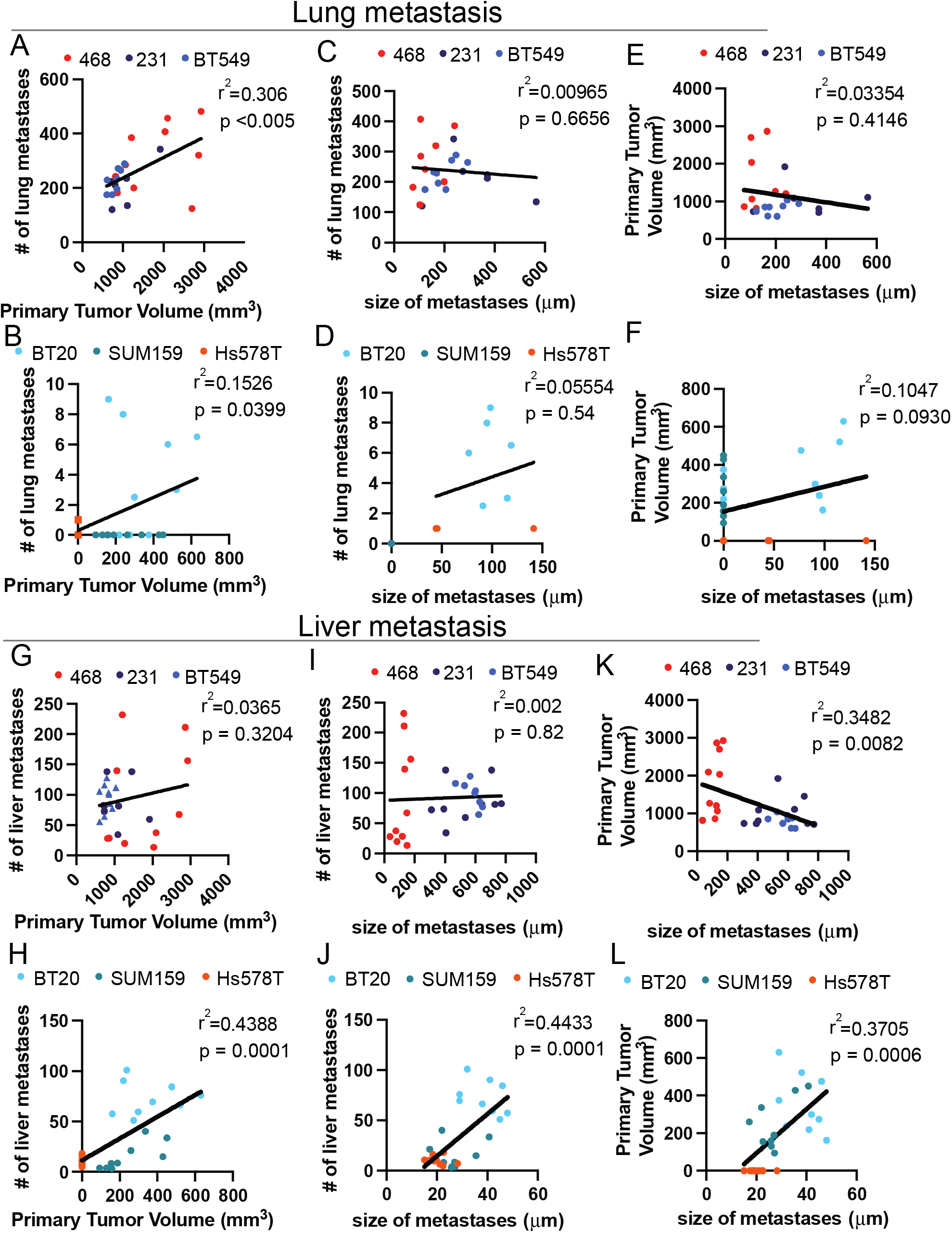
Correlation between tumorigenicity and metastasis in high and low metastatic potential cell lines. Correlation between high tumorigenic cell lines A) tumor volume and the number of lung metastasis B) tumor volume and size of lung metastasis C) size of lung metastasis and tumor volume. Correlation between intermediate and low tumorigenic cell lines D) tumor volume and the number of lung metastasis E) tumor volume and size of lung metastasis F) size of lung metastasis and tumor volume. Correlation between high metastatic potential cell line’s G) tumor volume and the number of liver metastasis H) tumor volume and size of liver metastasis I) size of liver metastasis and tumor volume. Correlation between low metastatic potential cell line’s J) tumor volume and the number of liver metastasis J) tumor volume and size of liver metastasis L) size of liver metastasis and tumor volume

**Supplemental Figure 3.**
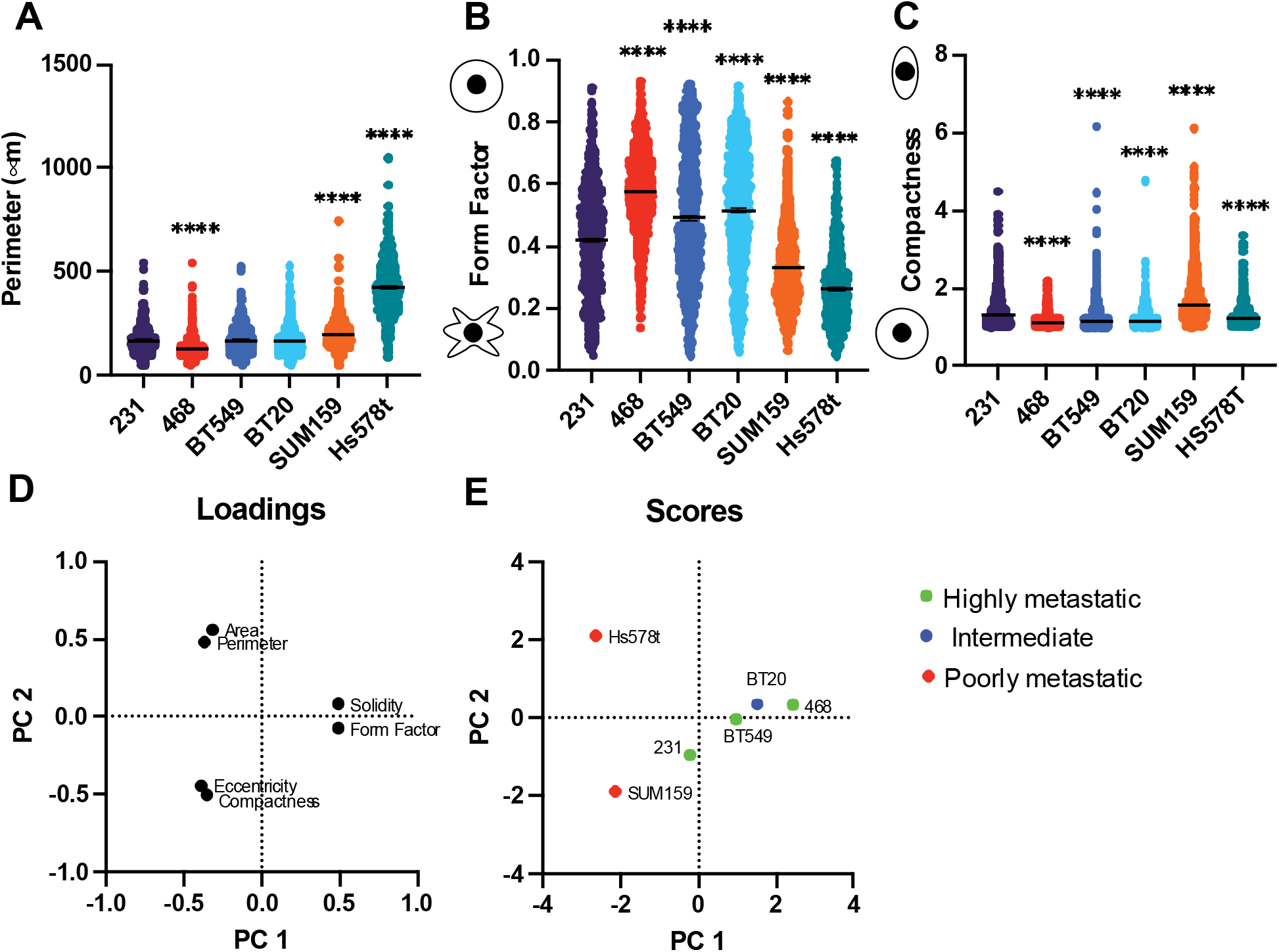
Characterization and clustering of cell line morphology. Quantification of cell shape parameters for each cell line 231 (n=679 cells), BT549 (n=799 cells), Hs578T (n=538 cells), and SUM159 (n=892 cells) (A) Perimeter (****P<0.0001) (B) Form Factor (****P<0.0001), (C) Compactness (****P<0.0001). Principal component analysis of the cell morphology metrics for each cell line (D) loadings scores of each cell morphology metric projected onto PC1 and PC2. (E) PC scores of each cell line projected onto PC1 and PC2 (E)Significance was determined using a one-way ANOVA with Dunnett’s multiple comparisons test compared to 231 controls. Each data point represents a single cell.

**Supplemental Figure 4.**
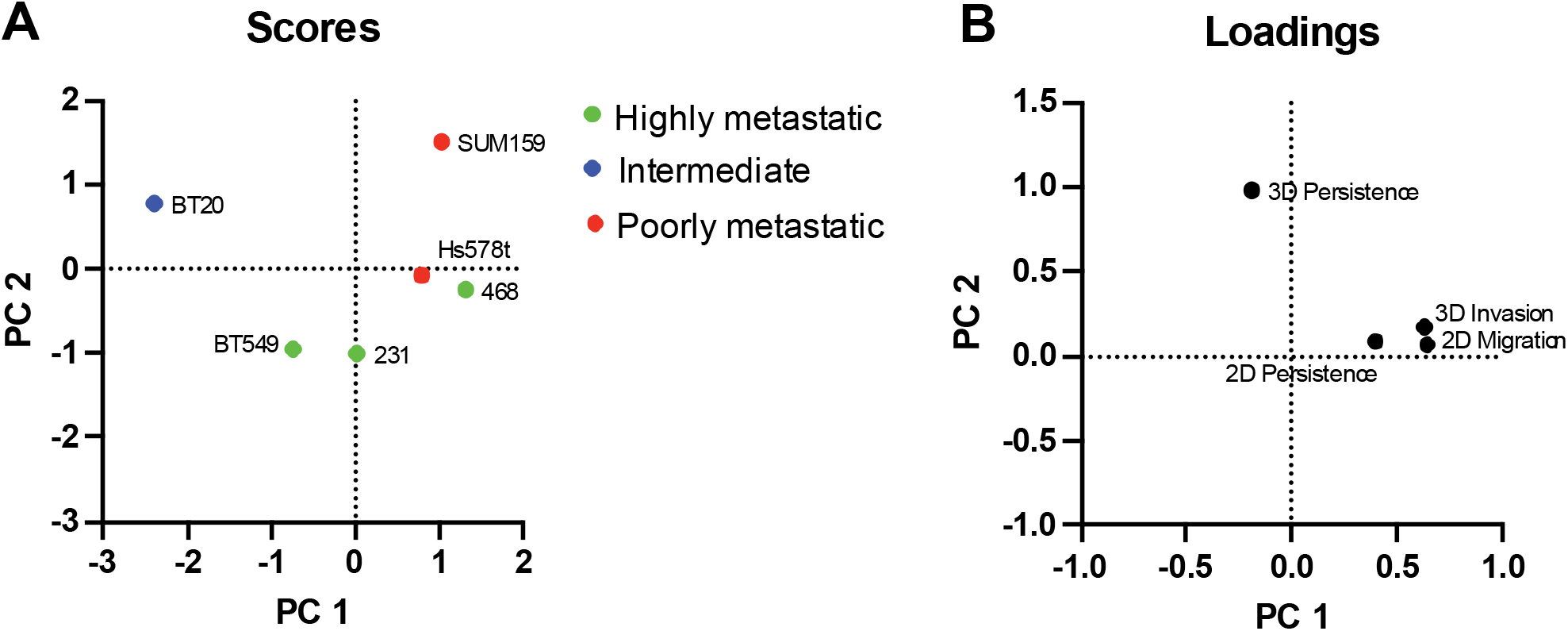
Characterization and clustering of cell lines 2D and 3D single cell motility. Principal component analysis of the single cell motility metrics for each cell line (A) PC scores of each cell line projected onto PC1 and PC2 (B) loadings scores of each cell morphology metric projected onto PC1 and PC2.

**Supplemental Figure 5.**
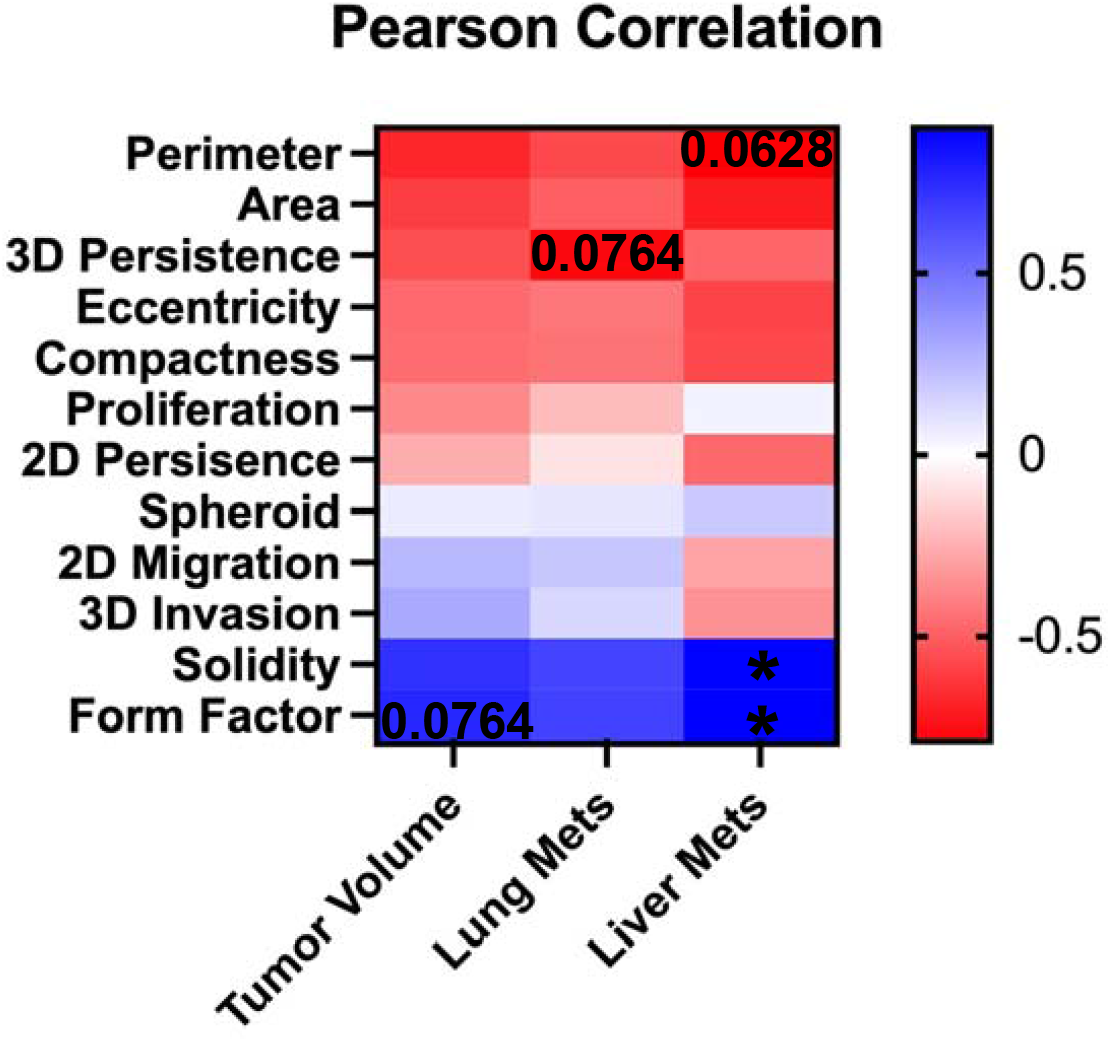
Spearman correlation between *in vitro* metrics and *in vivo* behavior.

## Notes

### Competing Interest Statement

The authors have declared no competing interest.

## References

[1] R. L. Siegel, K. D. Miller, H. E. Fuchs, and A. Jemal, “Cancer statistics, 2022,” CA. Cancer J. Clin., vol. 72, no. 1, pp. 7–33, 2022, doi: 10.3322/caac.21708.

[2] J. L. Moss, Z. Tatalovich, L. Zhu, C. Morgan, and K. A. Cronin, “Triple-negative breast cancer incidence in the United States: ecological correlations with area-level sociodemographics, healthcare, and health behaviors,” Breast Cancer, vol. 28, no. 1, pp. 82–91, Jan. 2021, doi: 10.1007/s12282-020-01132-w.

[3] D. Kaplan, “Overview of the Updated NCCN Guidelines on Triple-Negative Breast Cancer,” HMP Glob. Learn. Netw., Dec. 2021, Accessed: Feb. 05, 2023. [Online]. Available: https://www.hmpgloballearningnetwork.com/site/jcp/jcp-special-report/overview-updated-nccn-guidelines-triple-negative-breast-cancer

[4] G. Bianchini, J. M. Balko, I. A. Mayer, M. E. Sanders, and L. Gianni, “Triple-negative breast cancer: challenges and opportunities of a heterogeneous disease,” Nat. Rev. Clin. Oncol., vol. 13, no. 11, Art. no. 11, Nov. 2016, doi: 10.1038/nrclinonc.2016.66.

[5] C. K. Anders, T. M. Zagar, and L. A. Carey, “The Management of Early-Stage and Metastatic Triple-Negative Breast Cancer: A Review,” Hematol. Oncol. Clin. North Am., vol. 27, no. 4, pp. 737–749, Aug. 2013, doi: 10.1016/j.hoc.2013.05.003.

[6] S. Valastyan and R. A. Weinberg, “Tumor metastasis: molecular insights and evolving paradigms,” Cell, vol. 147, no. 2, pp. 275–292, Oct. 2011, doi: 10.1016/j.cell.2011.09.024.

[7] L. Ritsma et al., “Intravital Microscopy Through an Abdominal Imaging Window Reveals a Pre-Micrometastasis Stage During Liver Metastasis,” Sci. Transl. Med., vol. 4, no. 158, pp. 158ra145–158ra145, Oct. 2012, doi: 10.1126/scitranslmed.3004394.

[8] B. Waclaw, I. Bozic, M. E. Pittman, R. H. Hruban, B. Vogelstein, and M. A. Nowak, “A spatial model predicts that dispersal and cell turnover limit intratumour heterogeneity,” Nature, vol. 525, no. 7568, pp. 261–264, Sep. 2015, doi: 10.1038/nature14971.

[9] J. M. Tse et al., “Mechanical compression drives cancer cells toward invasive phenotype,” Proc. Natl. Acad. Sci., vol. 109, no. 3, pp. 911–916, Jan. 2012, doi: 10.1073/pnas.1118910109.

[10] P. Friedl and K. Wolf, “Tumour-cell invasion and migration: diversity and escape mechanisms,” Nat. Rev. Cancer, vol. 3, no. 5, Art. no. 5, May 2003, doi: 10.1038/nrc1075.

[11] M. J. Oudin and V. M. Weaver, “Physical and Chemical Gradients in the Tumor Microenvironment Regulate Tumor Cell Invasion, Migration, and Metastasis,” Cold Spring Harb. Symp. Quant. Biol., vol. 81, pp. 189–205, 2016, doi: 10.1101/sqb.2016.81.030817.

[12] A. J. Minn et al., “Distinct organ-specific metastatic potential of individual breast cancer cells and primary tumors,” J. Clin. Invest., vol. 115, no. 1, pp. 44–55, Jan. 2005, doi: 10.1172/JCI22320.

[13] A. Nguyen, M. Yoshida, H. Goodarzi, and S. F. Tavazoie, “Highly variable cancer subpopulations that exhibit enhanced transcriptome variability and metastatic fitness,” Nat. Commun., vol. 7, no. 1, Art. no. 1, May 2016, doi: 10.1038/ncomms11246.

[14] M. J. Oudin et al., “MENA Confers Resistance to Paclitaxel in Triple-Negative Breast Cancer,” Mol. Cancer Ther., vol. 16, no. 1, pp. 143–155, Jan. 2017, doi: 10.1158/1535-7163.MCT-16-0413.

[15] J. P. Fatherree, J. R. Guarin, R. A. McGinn, S. P. Naber, and M. J. Oudin, “Chemotherapy-induced collagen IV drives cancer cell invasion through activation of Src/FAK signaling.” bioRxiv, p. 2021.04.01.438074, Apr. 02, 2021. doi: 10.1101/2021.04.01.438074.

[16] J. R. Guarin, J. P. Fatherree, and M. J. Oudin, “Chemotherapy treatment induces pro-invasive changes in liver ECM composition,” Matrix Biol., vol. 112, pp. 20–38, Sep. 2022, doi: 10.1016/j.matbio.2022.08.002.

[17] K. J. Simpson et al., “Identification of genes that regulate epithelial cell migration using an siRNA screening approach,” Nat. Cell Biol., vol. 10, no. 9, Art. no. 9, Sep. 2008, doi: 10.1038/ncb1762.

[18] M. Lintz, A. Muñoz, and C. A. Reinhart-King, “The Mechanics of Single Cell and Collective Migration of Tumor Cells,” J. Biomech. Eng., vol. 139, no. 2, Jan. 2017, doi: 10.1115/1.4035121.

[19] W. J. Polacheck, I. K. Zervantonakis, and R. D. Kamm, “Tumor cell migration in complex microenvironments,” Cell. Mol. Life Sci. CMLS, vol. 70, no. 8, pp. 1335–1356, Apr. 2013, doi: 10.1007/s00018-012-1115-1.

[20] P.-H. Wu et al., “Single-cell morphology encodes metastatic potential,” Sci. Adv., vol. 6, no. 4, p. eaaw6938, Jan. 2020, doi: 10.1126/sciadv.aaw6938.

[21] P.-H. Wu et al., “Evolution of cellular morpho-phenotypes in cancer metastasis,” Sci. Rep., vol. 5, no. 1, Art. no. 1, Dec. 2015, doi: 10.1038/srep18437.

[22] J. P. Baskaran et al., “Cell shape, and not 2D migration, predicts extracellular matrix-driven 3D cell invasion in breast cancer,” APL Bioeng., vol. 4, no. 2, p. 026105, May 2020, doi: 10.1063/1.5143779.

[23] S. I. Fraley et al., “A distinctive role for focal adhesion proteins in three-dimensional cell motility,” Nat. Cell Biol., vol. 12, no. 6, Art. no. 6, Jun. 2010, doi: 10.1038/ncb2062.

[24] P.-H. Wu, S. S. Gambhir, C. M. Hale, W.-C. Chen, D. Wirtz, and B. R. Smith, “Particle tracking microrheology of cancer cells in living subjects,” Mater. Today, vol. 39, pp. 98–109, Oct. 2020, doi: 10.1016/j.mattod.2020.03.021.

[25] K. M. Charoen, B. Fallica, Y. L. Colson, M. H. Zaman, and M. W. Grinstaff, “Embedded multicellular spheroids as a biomimetic 3D cancer model for evaluating drug and drug-device combinations,” Biomaterials, vol. 35, no. 7, pp. 2264–2271, Feb. 2014, doi: 10.1016/j.biomaterials.2013.11.038.

[26] C. L. Yankaskas et al., “A microfluidic assay for the quantification of the metastatic propensity of breast cancer specimens,” Nat. Biomed. Eng., vol. 3, no. 6, Art. no. 6, Jun. 2019, doi: 10.1038/s41551-019-0400-9.

[27] E. Y. Lasfargues and L. Ozzello, “Cultivation of Human Breast Carcinomas2,” JNCI J. Natl. Cancer Inst., vol. 21, no. 6, pp. 1131–1147, Dec. 1958, doi: 10.1093/jnci/21.6.1131.

[28] X. Dai, H. Cheng, Z. Bai, and J. Li, “Breast Cancer Cell Line Classification and Its Relevance with Breast Tumor Subtyping,” J. Cancer, vol. 8, no. 16, pp. 3131–3141, Sep. 2017, doi: 10.7150/jca.18457.

[29] K. A. Janes and M. B. Yaffe, “Data-driven modelling of signal-transduction networks,” Nat. Rev. Mol. Cell Biol., vol. 7, no. 11, Art. no. 11, Nov. 2006, doi: 10.1038/nrm2041.

[30] Y. Yao, Y. Chu, B. Xu, Q. Hu, and Q. Song, “Risk factors for distant metastasis of patients with primary triple-negative breast cancer,” Biosci. Rep., vol. 39, no. 6, p. BSR20190288, Jun. 2019, doi: 10.1042/BSR20190288.

[31] M. Pistelli et al., “Prognostic Factors in Early-stage Triple-negative Breast Cancer: Lessons and Limits from Clinical Practice,” Anticancer Res., vol. 33, no. 6, pp. 2737–2742, Jun. 2013.

[32] A. Rizwan, M. Cheng, Z. M. Bhujwalla, B. Krishnamachary, L. Jiang, and K. Glunde, “Breast cancer cell adhesome and degradome interact to drive metastasis,” Npj Breast Cancer, vol. 1, no. 1, Art. no. 1, Oct. 2015, doi: 10.1038/npjbcancer.2015.17.

[33] A. J. Minn et al., “Genes that mediate breast cancer metastasis to lung,” Nature, vol. 436, no. 7050, pp. 518–524, Jul. 2005, doi: 10.1038/nature03799.

[34] X. Jin et al., “A metastasis map of human cancer cell lines,” Nature, vol. 588, no. 7837, Art. no. 7837, Dec. 2020, doi: 10.1038/s41586-020-2969-2.

[35] L. Gómez-Cuadrado, N. Tracey, R. Ma, B. Qian, and V. G. Brunton, “Mouse models of metastasis: progress and prospects,” Dis. Model. Mech., vol. 10, no. 9, pp. 1061–1074, Sep. 2017, doi: 10.1242/dmm.030403.

[36] A. Zaritsky et al., “Interpretable deep learning uncovers cellular properties in label-free live cell images that are predictive of highly metastatic melanoma,” Cell Syst., vol. 12, no. 7, pp. 733-747.e6, Jul. 2021, doi: 10.1016/j.cels.2021.05.003.

[37] S. M. Lyons et al., “Changes in cell shape are correlated with metastatic potential in murine and human osteosarcomas,” Biol. Open, vol. 5, no. 3, pp. 289–299, Feb. 2016, doi: 10.1242/bio.013409.

[38] A. L. Wishart et al., “Decellularized extracellular matrix scaffolds identify full-length collagen VI as a driver of breast cancer cell invasion in obesity and metastasis,” Sci. Adv., vol. 6, no. 43, p. eabc3175, Oct. 2020, doi: 10.1126/sciadv.abc3175.

[39] A. Naba, K. R. Clauser, J. M. Lamar, S. A. Carr, and R. O. Hynes, “Extracellular matrix signatures of human mammary carcinoma identify novel metastasis promoters,” eLife, vol. 3, p. e01308, Mar. 2014, doi: 10.7554/eLife.01308.

[40] B. R. Seo et al., “Obesity-dependent changes in interstitial ECM mechanics promote breast tumorigenesis,” Sci. Transl. Med., vol. 7, no. 301, p. 301ra130, Aug. 2015, doi: 10.1126/scitranslmed.3010467.

[41] L. Ling et al., “Obesity-Associated Adipose Stromal Cells Promote Breast Cancer Invasion through Direct Cell Contact and ECM Remodeling,” Adv. Funct. Mater., vol. 30, no. 48, p. 1910650, 2020, doi: 10.1002/adfm.201910650.

